# Gaming the cancer-immunity cycle by synchronizing the dose schedules

**DOI:** 10.1101/2024.10.31.621326

**Authors:** S. Mahmoodifar, P.K. Newton

## Abstract

We introduce a mathematical model of the cancer-immunity cycle and use it to test several hypotheses regarding the combination, timing, and optimization associated with chemotherapy and immunotherapy dosing schedules in the context of competition and selection pressure. A key conceptual idea is the value of synchronizing the dosing schedules with the fundamental period of the cancer-immunity cycle. The competitors in the population dynamics evolutionary game are the cancer cells, healthy cells, and T-cells, which form a non-transitive rock-paper-scissor chain, mediated by the tumor microenvironment. The chemotherapy and immunotherapy dosing schedules each act as control functions whose timing we synchronize with the fundamental period of the underlying nonlinear dynamical system. With the model, we show among other more detailed results, that chemotherapy and immunotherapy schedules are non-transitive; the best duration of the chemotherapy is around one-quarter of the cancer-immunity cycle, whereas for immunotherapy it is one-half cycle; immunotherapy dosing should preceed chemotherapy dosing. A general conclusion is that optimized timing of the dosing schedules can make up for lower total dose, opening up new possibilities for designing less toxic and more efficacious dosing regimens with drugs currently in use. Obtaining and calibrating more accurate measurements of the cycle-period across patient populations would be an important step in making some of these ideas clinically actionable.

## I. INTRODUCTION

Tumors never develop in isolation, they grow and metastasize in the context of their regulatory environment [1] which includes the tumor microenvironment supporting and regulating, among other things, our innate and adaptive immune system [2]. The highly coupled and symbiotic dynamics that ensues between a growing tumor and our active immune response can be significantly altered by systemic chemotherapy agents acting on the tumor cell population and immunotherapy treatments [3–10] acting via multiple mechanisms [6] on the T-cell population, both of which have the tremendous potential to drive the coupled tumor-immune system towards a less lethal state.

But that potential also carries with it the ability to trigger dangerous auto-immune inflammatory responses as well as negative impacts associated with high toxicities [13], and the evolution of resistance to therapy [14, 15]. As a result, there is a great need for rationally designed dosing protocols that are optimized to avoid some of these dangers but also to enable the many benefits that biotherapy can provide, if administered in conjunction the chemotherapy [16]. One way of doing this is to identify the inherent time-scales and periodicities of the underlying processes [17] and develop optimized dosing schedules that exploit them. This is in contrast to current standard-of-care dosing schedules, such as maximum-tolerated-dose or metronomic schedules [18], where the timing of the dosages are pre-scheduled, generally at fixed time intervals and fixed dosing levels.

The evolutionary game theory mathematical model of the cancer-immune cycle we develop allows us to explore the consequences of synchronizing dose schedules with the timescales and periodicities of the underlying system, both duration and sequencing, first with single and multi-pulsed dosing protocols, and then more complex dosing schedules designed using mathematical optimal control theory. A key and surprising result of the model is that dose timing can be more impactful than total dose delivered.

### A. The cancer-immunity cycle

A key timescale and periodicity associated with the immune system is driven by the complex, balanced, and regulated interplay between cancer cells, healthy cells, and T-cells, captured elegantly in a framework called the *cancer-immunity cycle* [11, 19], *a concise uni-directional seven-step cyclic chain of events highlighting its major steps. The first two steps are the release and presentation of cancer cell antigens, the key molecule over-expressed on the cancer cell surface that elicits T-cell activiation and allows the T-cells to distinguish the cancer cells from normal healthy cells. The remaining five steps all involve the ramping up and targeted attack on the tumor by the T-cell population. This includes priming and activation of the T-cells, trafficking to and infiltration into tumors, recognition of cancer cells by T-cells, and finally, killing of the cancer cells. Killing the cancer cells releases additional antigens as the cycle begins again, over and over, typically leading to an increase in accumulated antigen levels over time. The process is cyclic and usually robust. Because of its cyclic nature, one can define an inherent periodicity associated with the cycle (P*_*CIC*_), and with it, a frequency, which we call the *driving frequency* of the cycle. If all goes well as the cycles repeat, the increase in the immune response is kept in check, making the cycle self-sustaining. The period of the cycle, of course, is made up of the timescales of each of the seven steps in the cycle which can be highly variable depending on patient specific factors, cancer type and staging, immunogencity of the tumor, the tumor microenvironment [2, 12] and other factors. But as a rule of thumb, it is generally estimated that the process of antigen presentation by dendritic cells to T-cells occurs within hours to a few days after antigen uptake [2, 12], T-cell activation and T-cell migration and infiltration takes place over a few days to a few weeks [2, 12], and actual impact on tumor size (i.e. regression) can take weeks to months [2, 12], hence should be thought of as the rate-limiting step.

Another potentially relevant driving frequency in the chain is that of the cancer cell cycle [20] included in our figure 1 (roughly 12 to 18 hours as compared to 18 to 24 hours for normal cells), but exactly what role this accelerated timescale plays in the cancer-immunity cycle, aside from its role in tumor growth, is far less clear. For example, is its frequency somehow synchronized with the cancer-immune cycle via cell signaling [21]? Many more detailed nuanced and complete descriptions can be found in the original paper which first highlighted and isolated the steps in the cycle [19], updated more recently [11], as well as in [22, 23].

**FIG 1.**
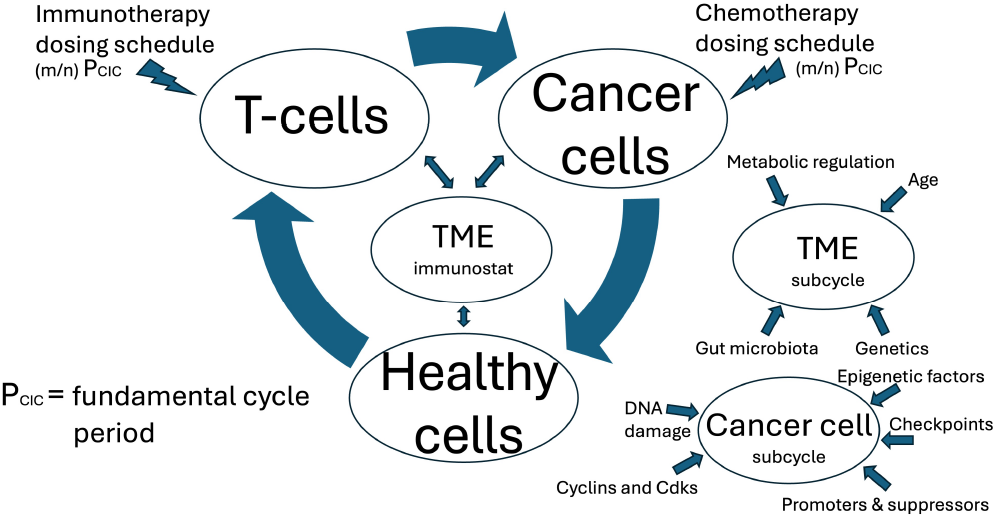
Key elements of the cancer-immunity cycle forming a non-transitive Cancer (rock) → Healthy (scissor) →T-cell (paper) chain. The tumor microenvironment (TME) acts as an immunostat, helping to regulate the immune response and partially determining the immunotype. Because of its importance, it can be thought of having its own subcycle [11]. The cancer cell cycle is also an important driver of the system. The cancer-immunity cycle has a fundamental period, *P*_*CIC*_, and frequency that are highly dependent of many factors. Our model develops the concept that the therapeutic frequencies should be applied in synchrony with a resonant frequency of the system, (*m/n*)*P*_*CIC*_, where *m* and *n* are integers.

Viewing the healthy cells, cancer cells, and T-cells as players in a population game, a useful framework in which to view their interaction chain is via analogy with the classic rock-paper-scissors game played by children in which rock crushes scissors, scissors cuts paper, and paper covers rock. In the cancer-immunity cycle, cancer cells outcompete healthy cells, healthy cells normally can keep the T-cells in check (homeostasis), yet T-cells have the ability to target and kill cancer cells. We detail this non-transitive chain and its relation to the cancer-immunity cycle in figure 1. But the rock-paper-scissor game [24], by itself, is too simple to capture the ecoevolutionary complexity of the true process, with its ability to adapt, select, and self-regulate.

The cancer-immunity cycle is initiated by the mutational accumulation of genetic alterations and a loss of normal cellular regulatory processes [19] which allows the tumor cell population to gain a selective advantage, accelerating its evolutionary fitness (measured in terms of reproductive prowess) relative to the surrounding healthy cell population. As a result, the tumor grows at the expense of the surrounding healthy tissue, while lowering the overall fitness of the entire collection of cells, rendering it vulnerable to chemotherapeutic agents, and because of its divergence from the normal cells, more easily recognized as foreign by the immune system [23]. As the tumor grows, the adaptive immune system is alerted and triggers a response involving T-cells, B-cells, NK-cells, macrophages, and a host of other cells [2] which attack the tumor cell population, slowing tumor growth, causing tumor regression. The three broad competing cell groups: healthy cells -tumor cells - T-cells, engage in a kind of evolutionary game, as depicted schematically in figure 1. In this cancer-immunity game, the T-cell population has the ability to alter the relative fitness of the tumor cell population (as compared to the fitness of the surrounding healthy cells), all via the tumor microenvironment (TME) which effectively functions as a regulatory immunostat [19]. The cancer cells, in turn, have the ability to elicit an immune response via the presentation and accumulation of the surface antigens which also serve as convenient targets for immunotherapies [25].

Going further and fleshing out more features of the cancer-immunity cycle and the role of the TME in both facilitating as well as suppressing the anti-cancer response (generally referred to as cancer immunoediting [26, 27]), it is well documented [28] that, if unregulated (either by the immune response or active therapeutic interventions) a tumor cell population grows in a Gompertzian-like [21, 28, 29] fashion, outcompeting the healthy cell population surrounding it. The activated immune response [11, 19] subsequently alters the nature of the ongoing competition between the healthy cells and cancer cells, effectively setting up the three-population evolving game. Viewing the immune system response as part of the tumor cell microenvironment, the regulatory environmental response functions as an immunostat [19] by lowering an overactive immune response and stimulating an underactive one. There is also abundant evidence that it acts as an important region in which metabolic regulation occurs [30], which has been suggested as a possible therapeutic target. In fact, because of the important role played by the TME, in particular its role in determining the *immunotype* of the tumor and its subsequent immune trajectory in a very dynamic process, the basic tumor-immunity cycle has been augmented in [11] to include what is called the tumor microenvironment cancer-immunity subcycle, which we include in figure 1. After regulation by the immune system, the cancer cell population can drop to a sufficiently low level, causing the immune response to settle back to its normal level associated with a homeostatic equilibrium [31], sometimes referred to as the cancer-immune set point (the details of which can depend on many factors including genetics, age, the microbiome — how much it varies from person to person is not well understood and is schematically represented as a generic bell curve of factors over the population in [23]), and the natural period and driving frequency of the cycle are established. Then, the cycle begins anew, the cancer cell population begins to regrow, which re-stimulates the immune response, etc. leading to an oscillatory exchange between the cancer cell population and the T-cell response with a time-lag based on individual physiology and other factors. Chemotherapy and immunotherapy agents administered exogeneously can, of course, further alter the nature of this cycle.

### B. A focus on timing

A key conceptual idea in this paper is that the chemotherapy and immunotherapy dosing schedules have their own periodicities and timing associated with their application, and that these should be synchronized with the driving frequency of the underlying cancer-immunity cycle in order to maximize their effect. In the physics and engineering literature, this concept is referred to as supplying external driving at a resonance frequency of the fundamental period of the oscillator [32] and is known to optimize the gain achieved from the external driving mechanism (e.g. pumping a swing at its natural frequency).

As far as we are aware, this concept of resonant forcing has not been exploited in biological contexts, although we draw attention to an interesting literature [33–35] which covers related ideas in the context of ecological and epidemiological systems, as in, for example, the synchronization of incidences of infectious diseases and seasonal fluctuations, and pulse vaccination efforts. In general terms, as discussed in [33, 35], one might anticipate the existence of a hierarchy of resonant frequencies of the external forcing functions (in our case, the chemotherapy and immunotherapy dosing schedules) synchronized with rational multiples (i.e (*m/n*)*P*_*CIC*_, where *m* and *n* are integers) of the natural period of oscillation of the system. In practice, because of the inherent complexities associated with precise measurments of the biomarkers necessary to measure long and complex periodicities (as well as the nonlinearities in the model system which generically makes resonant curves amplitude dependent), we focus on small values of *m* and *n*: *m* = 1, *n* = 1, 2, 4.

More generally, as pointed out in the review paper [17], the concept of focusing on the timing and coordination of chemotherapy and immunotherapy treatments in order to increase efficacy and reduce toxicity is only at the very beginning stages of being understood and implemented with the need for rational combination treatment schedule design to replace empirical guess and test type of approaches. Their figure 1 [17] nicely summarizes much of what is known through carefully designed clinical trials on temporally programmed combination cancer immunotherapies in relation to the seven steps in the cancer-immunity cycle. Our goal in developing and testing the mathematical model described in the following sections is to build a mathematical and computational framework to begin to quantitatively test several hypotheses associated with some of these ideas.

### C. Evolutionary games and mathematical control theory

To understand this better, we introduce an evolutionary game for the cancer-immunity cycle, played among the healthy cell population (*H*), the cancer cell population (*S* denoting chemosensitive tumor cells), and the T-cell population (*n*), with additional external control functions that model the chemotherapy (*C*(*t*)) dosing schedule and the immunotherapy (*θ*(*t*)) dosing schedule. Our goal is to develop a simple interpretable model that focuses on the symbiotic and co-evolving dynamics and competition that ensues between (*H, S, n*), framed as an evolutionary game with the addition of the tools of nonlinear control theory [36, 37] with controllers 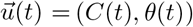.To simplify our model from the original seven steps of the cancer-immunity chain [11, 19], we do not explicitly model the details of each step, but, for example, lump together steps 3-5 carried out by the immune-system components into one compartmental variable, *n*(*t*), representing the induced T-cell response. Likewise, the detailed processes carried out by the steps involving the cancer cells are encoded in a fitness payoff matrix which determines the dynamics of the cancer cell variable, *S*, through their competition with the healthy cell variable *H*, each normalized so they are represented as proportions (fractions) in the total population of cells we consider. We then couple these fractional equations to a volumetric tumor growth law for total volume *V* (*t*). Although we are not aware of any specific mathematical model of the cancer-immunity cycle, there have been many previous approaches developing more detailed compartment models for the tumor-immune system, along with control theory, such as [38–41] and the reviews [42, 43], accounting for a more fine-scaled deliniation of species, with multiple parameters that can be fit to patient data and used for forecasting. These models have offered many important insights and continue to play a key role in developing a quantitative and predictive understanding of the important biological processes at play.

### D. What questions does our model address?

Our goal in framing the problem as an evolutionary game for the cancer-immunity cycle is to highlight key features of the trade-offs and the nature of the competition the three cell populations confront, highlight the importance of establishing the fundamental period of the cycle, and focus on the value of synchronizing therapeutic dose schedules with rational multiples of this fundamental period. By doing this, we then can address several basic questions regarding combination immunochemotherapy [44] in the context of a co-evolving system subject to selection pressure. These questions include:

Q1. Are the chemotherapy and immunotherapy schedules transitive, i.e. do they have the same effect if the order in which they are applied is reversed? Answering this might motivate further more detailed studies on optimal sequencing of therapeutic dose schedules [45].

Q2. How to best combine the different chemotherapy and immunotherapy dose schedules, sequentially or concomitantly [46, 47]? Answering this might lead to more direct and detailed studies comparing these two options in murine models or targeted clinical trials to shed light on a currently unresolved question [16, 44].

Q3. When is the optimal time to start and stop the immunotherapy? Answering this unresolved questions could motivate further studies to better understand this in different contexts [48] which could lead to insights into optimal timing of adaptive schedules.

Q4. What is the benefit of using more complicated time-dependent dosing schedules? While currently impractical in clinical practice because of the challenges associated with real-time biomarker tracking of cell populations, demonstrating the benefit of adaptive control theory procedures could spur the development of better and less invasive monitoring procedures based, for example, on blood-based tumor genotyping and other molecular diagnostic strategies that allow the possibility of bringing *time*, as an additional variable, into the arsenal of precision medicine [49].

These and other questions will be addressed in what follows in the context of our evolutionary game-theory model.

## II. MODEL ELEMENTS

The model we develop is based on that proposed in in a different context in which environmental effects alter the nature of the payoffs in an evolutionary game [51, 52]. The authors call these *feedback evolving* games, or *co-evolutionary* game theory. They extend the form of the replicator dynamical system [51, 52] to include dynamical changes to the environment and then use their model to revisit the tragedy of the commons scenario, in which maximizing individual payoffs leads to an overall payoff deficit, to address what happens if overexploitation of a resource changes incentives for further action. Our model can be thought of similarly, where we treat the healthy cell and the cancer cell populations (measured as fractions in a deterministic set-up as is typically done within the evolutionary game theory framework) as an evolving competition with a nominal prisoner’s dilemma (PD) payoff matrix with healthy cells playing the role of cooperators, and cancer cells playing the role of defectors [37, 53–56]. With this choice, the cancer cell population has higher fitness (growth rates) than the healthy cell population in the absence of therapeutic intervention, making them our therapeutic target. The immune system dynamics (T-cell level) in our model is treated as an environmental response that can adjust the payoff matrix entries governing the healthy-cancer cell interactions, altering the game and indirectly lowering the cancer cell fitness, as we describe in more detail next.

### A. Model description

Denoting the healthy cell (*H*), cancer cell (*S*), immune system response by the variables (*x*_1_(*t*), *x*_2_(*t*), *n*(*t*)), with *x*_1_ + *x*_2_ = 1 (0 ≤*x*_1_ ≤1; 0 ≤*x*_2_ ≤1, 0 ≤*n* ≤1), our model is based on the two-component replicator dynamical system [52] governing the healthy/cancer cell fractions (*x*_1_(*t*), *x*_2_(*t*)), coupled to a logistic type of equation measuring the immune system response *n*(*t*):[eq

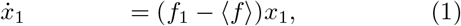

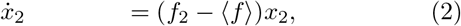

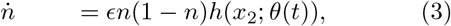

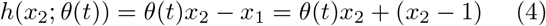

The equations are supplemented with initial conditions (*x*_1_(0), *x*_2_(0), *n*(0)), with *n*(0)∈ [0, 1]; *x*_1_(0)∈ [0, 1]; *x*_2_(0)∈ [0, 1]; *x*_1_(0) + *x*_2_(0) = 1, and a 2*×* 2 payoff matrix *A*(*n*) (coupled to the immune response) defining the fitness profiles, *f*_1_ and *f*_2_, of each sub-population, and the population average fitness ⟨*⟨f⟩*.

The function *h*(*x*_2_; *θ*(*t*)) is the instanteous growth rate of the T-cell population (*n*(*t*)), with *θ*(*t*) playing the role of our immunotherapy control function, with *θ*∈ [2, 5]. The baseline immune-system level (no immunotherapy) is set at *θ* = 2, while when *θ >* 2, immunotherapy is turned on, indirectly applying selection pressure on the healthy cell population by lowering the threshold value of *x*_2_ at which *h >* 0. When *h >* 0, the T-cell population level rises and *n*→ 1 (a fixed point), and when *h <* 0, it decays and *n* →0 (also a fixed point). In these equations, *f*_1_ represents the fitness of the *x*_1_ population, *f*_2_ represents the fitness of the *x*_2_ population, and ⟨*⟨f⟩* represents the population average fitness. A key variable in our system is *f*_2_ −⟨*⟨f⟩* ≡*α*_*G*_, which is the instantaneous growth rate of the cancer cell fraction. The fitness functions are defined as:

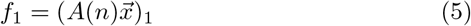

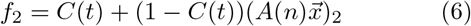

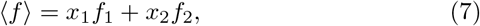

with *C*(*t*) ∈ [0, 1] playing the role of our chemotherapy control function. The subscript in eqn (5) indicates that it is the first component of the vector 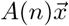,while the subscript in eqn (6) indicates that it is the second component. When *C*≡ 0, there is no chemotherapy acting on the cancer cells and the payoff matrix determines selection, when *C >* 0, chemotherapy is turned on and applies selection pressure on the healthy cell population.

Because the model is based on tumor cell/healthy cell *fractions*, we need an additional equation to track tumor volume, *V* (*t*), as the cancer cell population grows and decays. For this, we use a logistic equation with growth rate *δ* + *α*_*G*_ and carrying capacity *K >* 1:

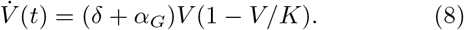

The (small) parameter *δ >* 0 models the inability of the natural immune reponse to completely counteract the overall fitness advantage the cancer cells have established over the healthy cells inducing tumor growth. While the immune response can slow growth and even cause (temporary) tumor regression, it generally cannot entirely eradicate the tumor cell population, particularly as the tumor gets large. As a result, without the additional benefit of immunotherapy and chemotherapy, the tumor will continue to grow, although generally not at a constant rate in a monotonic fashion. A representative volumetric growth curve for the tumor during six cancer-immunity cycles is shown in figure 9. This will be discussed at greater length later.

The relevance of the replicator eqns (1), (2) as the governing equations for (well-mixed) natural selection dynamics governing the healthy/cancer cell competition is: (i) the growth rate of each sub-population is proportional to its fitness (relative to the population average); (ii) fitness is population-dependent [57]; and (iii) there is a mechanism of replication of sub-populations [52] favoring the one that has higher fractional representation in the population. This is discussed in more detail in [37, 52–55].

### B. The governing payoff matrix

The 2*×* 2 payoff matrix *A*(*n*) determining fitness has entries that are functions of the T-cell response level *n*(*t*) (0≤ *n*≤ 1), which we take as an overall measure of immune-system activity (i.e. it involves many components and cell types that we do not model individually). *ϵ* is a parameter (0≤ *ϵ*≤ 1) that determines the relative timescales between the immune response and the cancer cell dynamics and controls the fundamental period of the cancer-immunity cycle (see figure 6 discussed later).

As the immune response rises, the tumor regresses; as the immune response drops, the tumor grows. Because of this, our payoff matrix *A* is constructed as a linear combination of two components, *A*_*G*_ and *A*_*R*_ that are responsible for tumor growth and tumor regression respectively, with a smooth function, *g*(*n*(*t*); *a*) that interpolates the two. *A* is defined by:

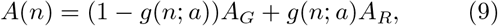

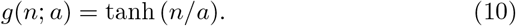

The two components of the payoff matrix *A* in eqn (9) are responsible for tumor growth, and tumor regression, respectively. Each is defined by:

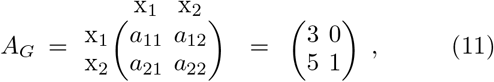

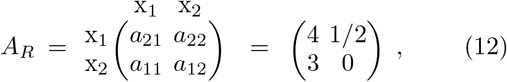

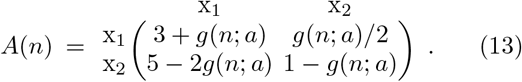

*A*_*G*_ is a standard prisoner’s dilemma matrix [52] since *a*_21_ *> a*_11_ *> a*_22_ *> a*_12_. The healthy cell population (*x*_1_) plays the role of cooperators, and cancer cell population (*x*_2_) plays the role of defectors, with defectors (*x*_2_ = 1) being the Nash equilibrium and an asymptotically stable equilibrium. In eqns (1), (2), using the tumor growth payoff matrix, *A*_*G*_, *a*_21_ − *a*_11_ = 2 *>* 0 is the (linear) growth rate of *x*_2_, relevant when *x*_2_ is small and *a*_12_ − *a*_22_ = − 1 *<* 0 is the linear decay rate of *x*_1_, relevant when *x*_2_ is large. This leads to tumor cell saturation, *x*_2_→ 1. Figure 2(a),(b) shows the nonlinear evolution to the Nash equilibrium *x*_2_ = 1 in this case. On the other hand, when the tumor regression payoff matrix, *A*_*R*_, is used, *a*_22_ − *a*_12_ = 1*/*2 *>* 0 is the linear growth rate of *x*_1_ relevant when *x*_1_ is small and *a*_11_− *a*_21_ = − 1 *<* 0 is the linear decay rate of *x*_2_ relevant when *x*_1_ is large. This leads to tumor cell extinction, *x*_2_ →0. Figure 2(c),(d) shows the nonlinear evolution to the Nash equilibrium *x*_2_ = 0 in this case. Notice that these choices of *A*_*G*_ and *A*_*R*_ reflect the fact that the immune system’s ability to reduce the fitness of the tumor cell population in the regression phase does not symmetrically compensate for the fitness advantage of the tumor cells over the healthy cells in the growth phase — growth and regression are not equal and opposite; the regression phase induced by the natural immune response is more gradual and takes longer than the more explosive growth phase induced by the mutational fitness advantage of the cancer cell population over the healthy cell population.

**FIG 2.**
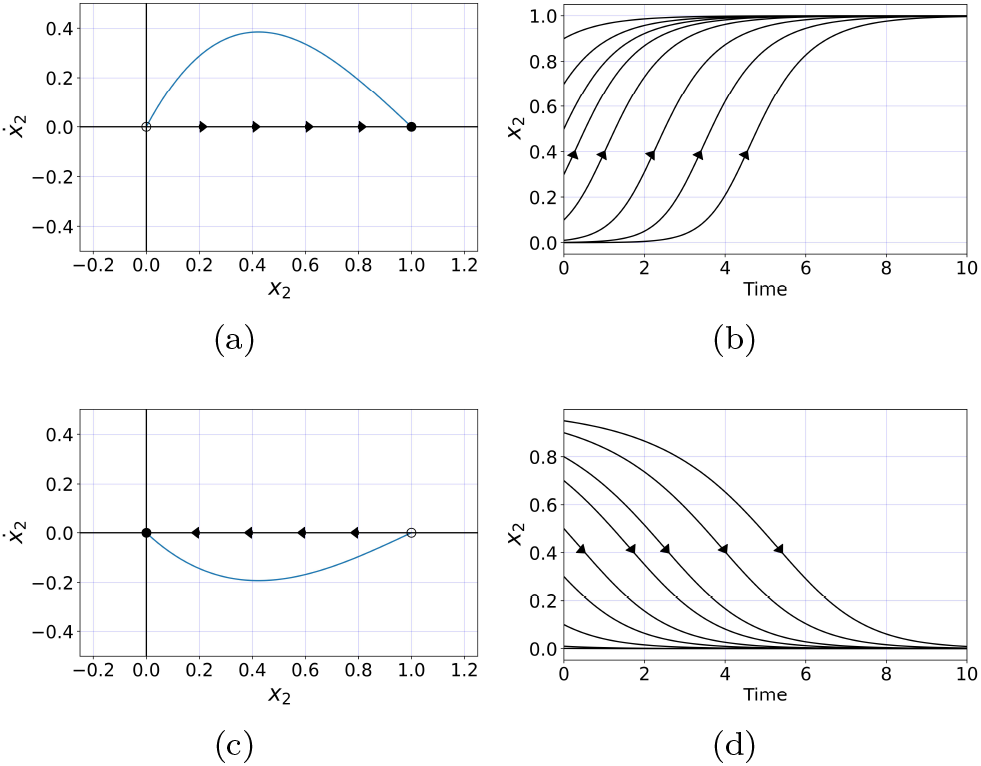
Tumor growth and tumor regression are determined by each of the two payoff matrices *A*_*G*_ and *A*_*R*_ respectively. Phase-portrait corresponding to the *A*_*G*_ payoff matrix of PD type modeling tumor growth. All defectors (*x*_2_ →1) is the ESS; (b) Tumor growth curve (S-shaped) associated with the *A*_*G*_ payoff matrix, *x*_2_→ 1; (c) Phase-portrait corresponding to the *A*_*R*_ payoff matrix, modeling tumor regression. Regression is slower than growth, reflecting the fact that the immune system’s ability to reduce the fitness of the tumor cell population in the regression phase does not symmetrically compensate for the fitness advantage of the tumor cells over the healthy cells in the growth phase. All cooperators (*x*_2_ → 0) is the ESS; (d) Tumor regression curve associated with *A*_*R*_, *x*_2_ → 0. This payoff matrix becomes increasingly dominant as the T-cell response variable *n*(*t*) → 1.

The interpolation function, *g*(*n*; *a*), is shown in figure 3(a) for parameter values *a* = 0.1, 0.2, 1, 2, 5. Since *g*(0; *a*) = 0, when there is no immune system response, *A*(0) ≡ *A*_*G*_ and the tumor growth is dictated by the growth payoff matrix (11). Since *g*^*/*^(0; *a*) = 1*/a*, small values of the parameter *a* gives rise to large slopes, indicating that the payoff matrix shifts quickly from *A*_*G*_ to a mixture of growth and regression as the T-cell level rises. In this sense, the parameter measures how sensitive the tumor cell response is to the initial rise in T-cell levels. As *n* →1, *g*(*n*; *a*) → tanh(1*/a*). In this case, the two coefficients (1 − tanh(1*/a*)) and tanh(1*/a*) weight the specific add-mixture of *A*_*G*_ and *A*_*R*_. We take *a* = 0.1 for our simulations. It should be noted that the fixed points of the system (1), (2), *x*_1_ = 0, *x*_2_ = 1 and *x*_1_ = 1, *x*_2_ = 0 are never reached in finite-time (i.e. the tumor never saturates or enters into complete remission as *x*_2_ *>* 0). Likewise, the fixed points of eqn (3), *n* = 0, 1 are never reached.

**FIG 3.**
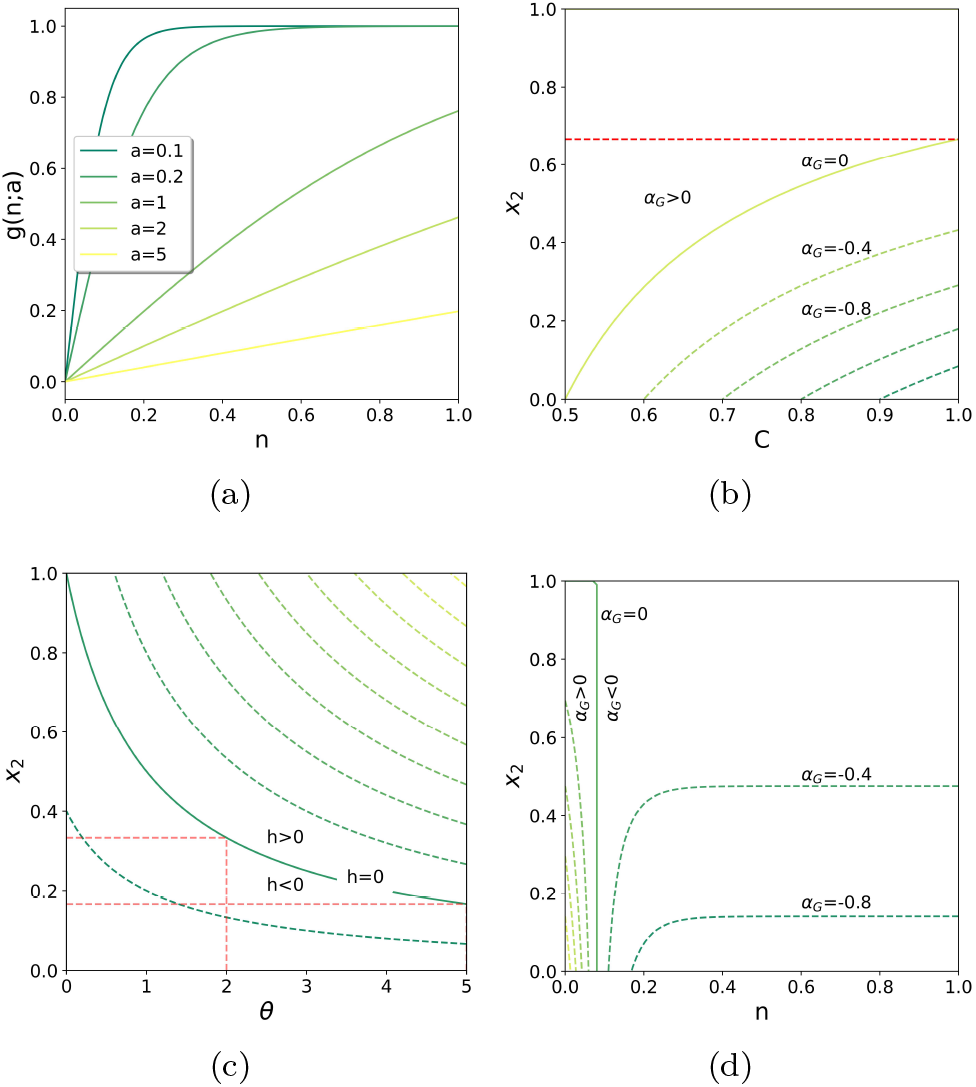
Further model elements. (a) Payoff matrix interpolation function *g*(*n*; *a*) = tanh(*n/a*) as a function of T-cell response *n* for different values of the response parameter *a*; Tumor cell growth rate level curves *α*_*G*_≡ *f*_2_ −⟨*f⟨*. in the (*C, x*_2_) plane with *n* = 0. The critical curve across which the growth rate changes sign is *α*_*G*_ = 0. (c) T-cell growth rate level curves, *h*(*x*_2_; *θ*) in the (*θ, x*_2_) plane. The critical curve on which the growth rate switches sign is *h*(*θ, x*_2_) = 0. (d) Tumor cell growth rate level curves *α*_*G*_≡ *f*_2_ −⟨*f⟨* in the (*n, x*_2_) plane with *C* = 0. The critical curve across which the growth rate changes sign is *α*_*G*_ = 0.

### C. Growth rates

To understand the role of the chemotherapy parameter *C*, the immunotherapy parameter *θ*, and the immune system level *n* in determining growth or decay of the tumor cell fraction, we focus on the growth rate functions *α*_*G*_(*x*_2_, *n*; *C*), and *h*(*x*_2_; *θ*), where:

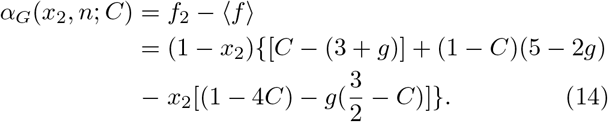

For the special case *n* = 0 (immuno-deficient), since *g*(0; *a*) = 0, the growth rate formula simplifies to:

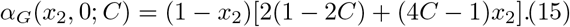

As shown in figure 3(b), when *C <* 1*/*2, *α*_*G*_ *>* 0 and the tumor cell population grows. When *C >* 1*/*2, if *x*_2_ is small enough to lie below the curve *α*_*G*_ = 0, *α*_*G*_ *<* 0 and the tumor cell population will regress. If *x*_2_ lies above the curve, the chemotherapy dose is not strong enough to cause regression, but will only slow the tumor growth. At its maximum, *C* = 1, the critical value at which *α*_*G*_ changes sign is *x*_2_ = 2*/*3. For tumor cell fractions above that value (with no immune response), no amount of chemotherapy can lead to tumor regression.

With an immune response (*n >* 0), the function *h*(*x*_2_; *θ*) in eqn (4) provides further important coupling of the cancer cell population to the immune system response since it plays the role of the instantaneous growth rate of the T-cell level. It is straightforward to see from eqn (4) that *h* switches sign when *θ* = *x*_1_*/x*_2_ = (1− *x*_2_)*/x*_2_. The critical curve *h*(*x*_2_; *θ*) = 0 is shown in figure 3(c) in the (*θ, x*_2_) plane. When the cancer cell fraction is sufficiently small, *h <* 0, which implies that the T-cell level decays towards its equilibrium value *n* = 0. As *x*_2_ increases, it crosses the critical curve so that *h >* 0, eliciting an immune system response. In this regime, the T-cell level begins to increase and the tumor regression component of the payoff matrix, *A*_*R*_, plays a bigger role, causing the tumor to regress. For *θ* = 2, which is our baseline with no immunotherapy, *h* switches from negative to positive when *x*_2_ exceeds 1*/*3. For *θ* = 5, when immunotherapy is on, *h* switches from negative to positive when *x*_2_ exceeds 1*/*6. As the tumor regresses, eventually *x*_2_ decreases so that the critical curve is crossed again, *h* switches from positive back to negative, and the immune response begins to diminish for the cycle to begin again. Finally, in figure 3(d) we show the growth rate *α*_*G*_ in the (*x*_2_, *n*) plane, with *C* = 0 which is instructive. For sufficiently small values of *n* (roughly *n <* 0.1), *α*_*G*_ *>* 0, and the immune system alone cannot induce tumor regression. For larger values (*n >* 0.1), *α*_*G*_ switches sign to negative, and the immune system begins to lower the tumor cell fraction *x*_2_.

Combining the two effects of chemotherapy and the immune response leads to better control of the tumor cell population. We show level curves of the growth rates *α*_*G*_ in the (*C, x*_2_) for different values of *n* in figure 4. Figure 4(a) first shows the threshold effect for *C* = 1*/*2 when there is no immune response. In order for *C* to cause tumor regression for even infinitesimal fractions of *x*_2_, *C* must be larger than 1*/*2. In figures 4(b),(c),(d) we show the critical level curve *α*_*G*_ = 0 shifting to the left as *n* increases -with the immune system acting together with chemotherapy, the critical value required for *C* to cause regression is lowered.

**FIG 4.**
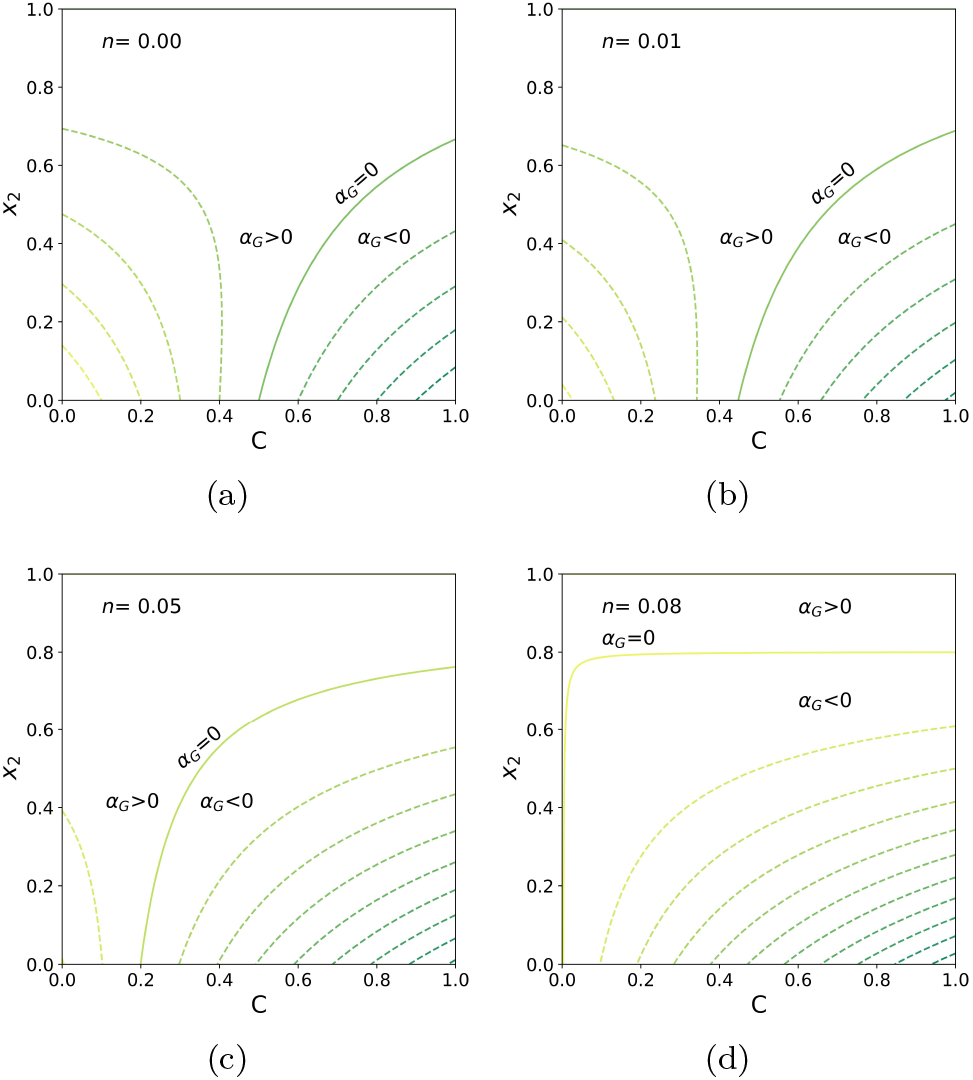
Tumor cell growth rate level curves *α*_*G*_≡ *f*_2_− ⟨*⟨f⟩* in the (*C, x*_2_) plane. The critical curve across which the growth rate changes sign is *α*_*G*_ = 0. (a) *n* = 0; (b) *n* = 0.01; (c) *n* = 0.05; (d) *n* = 0.08.

Details of the system response to our chemotherapy parameter *C* depend, of course, on the initial conditions in the model. Figure 5 shows the combined effects of chemotherapy and the immune system level for different initial values the the tumor cell fraction *x*_2_ in a sequence of simulations for fixed values *C* and *n*. Figures 5(a),(b),(c) shows the time evolution of *x*_2_(*t*) for *C* = 0.4, 0.6, 1.0, in the immuno-deficient case *n* = 0. Below the threshold *C* = 1*/*2, *x*_2_→ 1 for all initial values. For *C* = 0.6 (above threshold), tumor regression occurs for small initial tumor cell fractions, but growth occurs for larger values. Even for values *C* = 1.0, figure 5(c) shows tumor growth ensues for sufficiently large inital tumor cell fractions. In figures 5(d),(e),(f), we keep *C* = 0, and show how larger values of the immune system level helps lower the growth of tumor cell populations, even for large initial fractions when *n* = 0.20 and higher. Figures 5(g),(h),(i) show the combined ability of chemotherapy and the immune system to lead to tumor regression. For example, figure 5(i) shows that with *C* = 0.7 and *n* = 0.05, regression occurs for initial tumor fractions *x*_2_ = 0.6, as compared with figure 5(e) which shows tumor growth for that same initial value, when there is no chemotherapy. It is these combined effects of chemotherapy and the immune system response, in a dynamical setting, that we examine in the next section.

**FIG 5.**
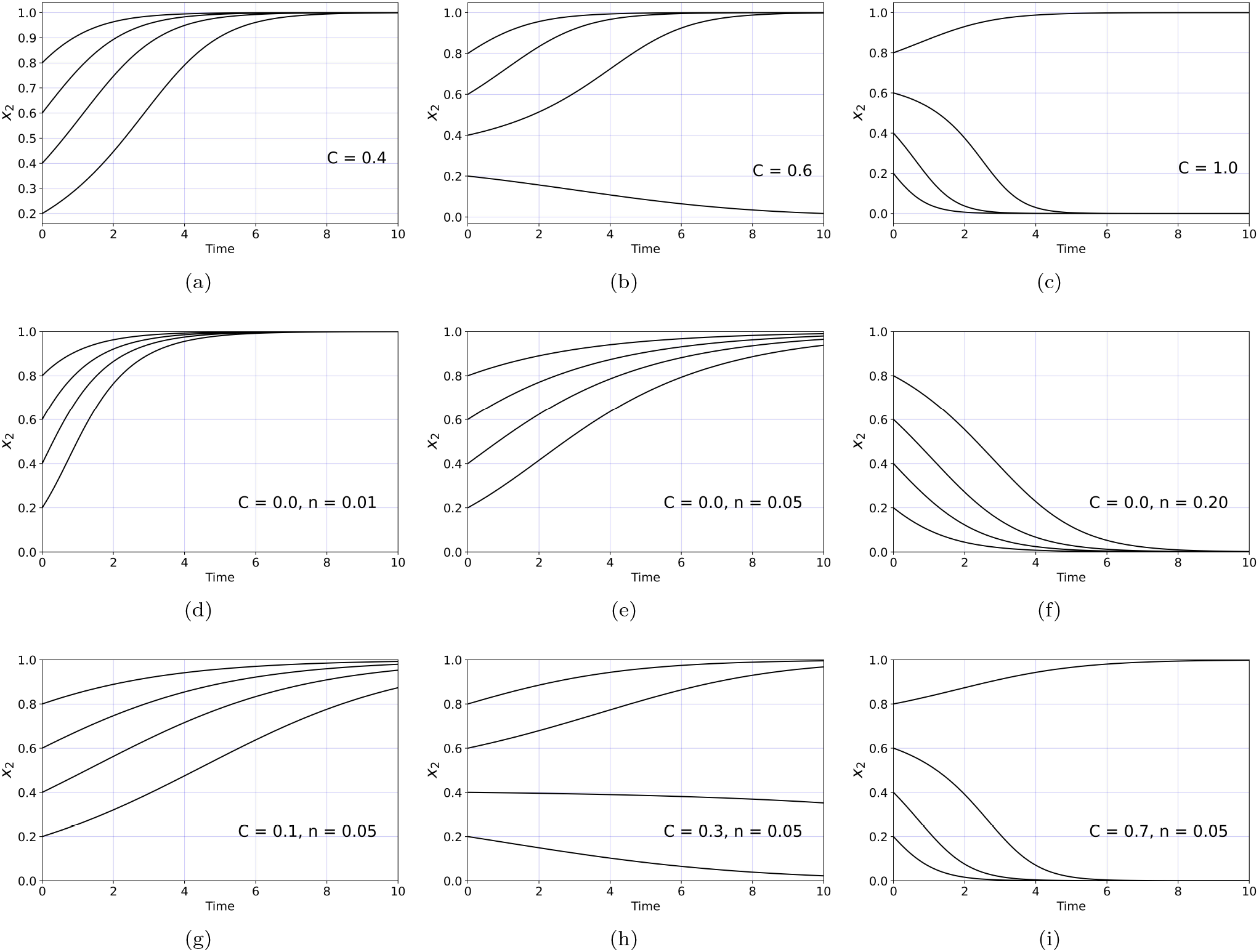
Tumor cell fraction as a function of *C* and *n*. (a) Immunodeficient *n* = 0, *C* = 0.4 (below threshold); (b) Immunodeficient *n* = 0, *C* = 0.6 (above threshold); (c) Immunodeficient *n* = 0, *C* = 1.0 (above threshold); (d) *C* = 0, *n* = 0.01 (e) *C* = 0, *n* = 0.05; (f) *C* = 0, *n* = 0.2;; (g) *C* = 0.1, *n* = 0.05; (h) *C* = 0.3, *n* = 0.05; (i) *C* = 0.7, *n* = 0.05.

**FIG 6.**
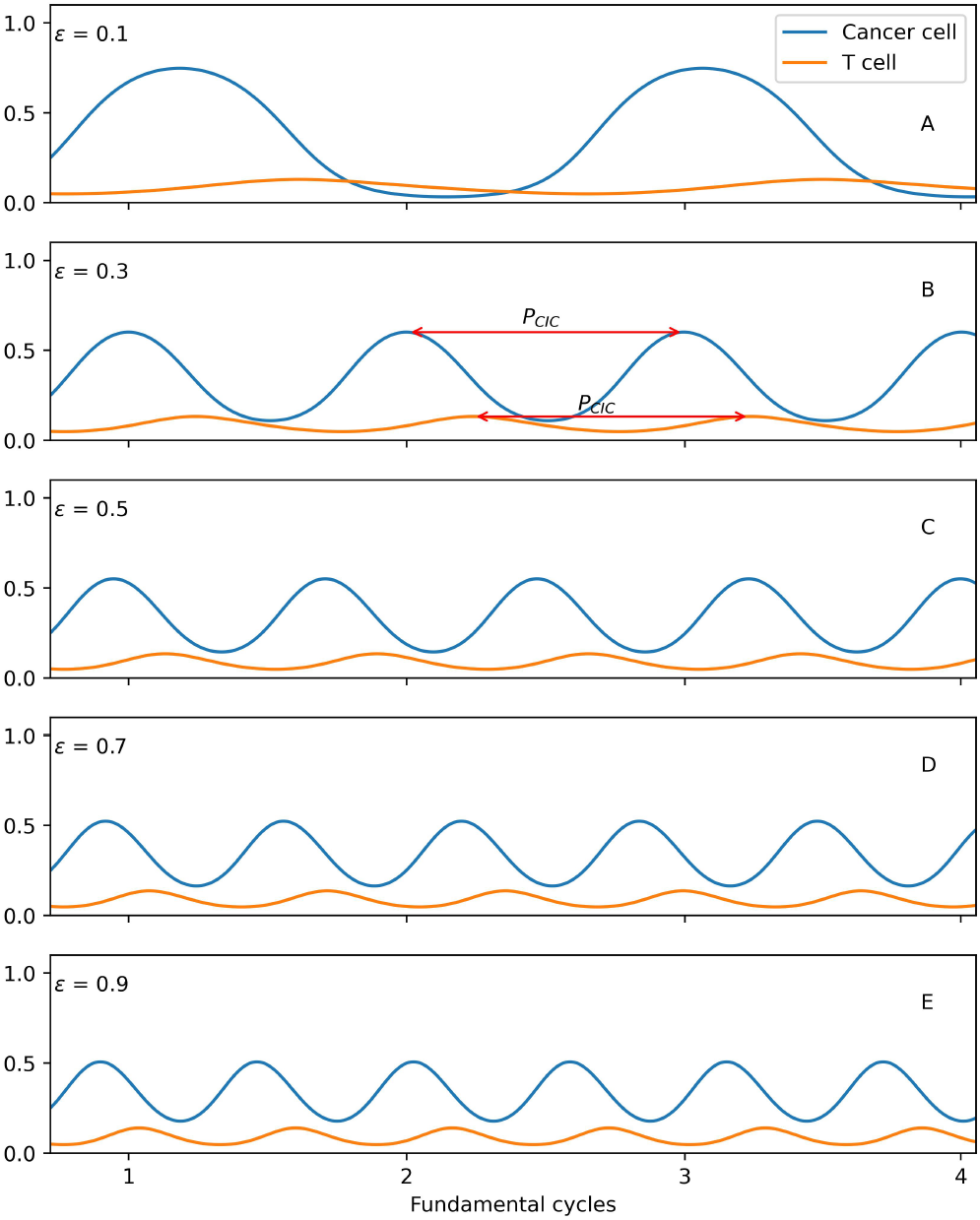
The fundamental cycle period and frequency scale with *c* with a phase shift between the cancer cell growth/regression phases and the immune system response. Time units are marked in cycle periods for ϵ = 0.3. A. ϵ = 0.1; B. ϵ = 0.3; C. ϵ = 0.5; D. ϵ = 0.7; E. ϵ = 0.9.

We make two final important points regarding the model. First, the model is dimensionless, whose parameters have also been rendered dimensionless. As such, the time variable in our model is measured in dimensionless time-units, so we show time progression in terms of the number of cycles marking the tumor cell population peaks. Establishing precise connections between the time variable and dimensional time would require more clinical data estimating, for example, the fundamental timeperiod and frequency of the underlying cancer-immunity cycle. Some of the details of these issues are discussed in [17]. A second point is that our model is deterministic, whereas one usually thinks of the cancer-immune problem as stochastic. One should think of our model as governing the expectation values associated with an underlying stochastic process which is nicely discussed in [58] in the context of co-evolutinary populations.

### D. Establishing the fundamental period of the cancer-immune cycle

In order to demonstrate some basic features of our model, we show some simple scenarios in figures 6-9 for ranges of relevant parameter values. Figure 6 shows the output of simulations of the coupled cancer cell/T-cell system for different values of the timescale parameter *ϵ* = 0.1, 0.3, 0.5, 0.7,, 0.9, the main parameter that establishes the fundamental period of the cancer-immunity cycle. For small values of *ϵ* (say *ϵ* = 0.1 shown in panel A), the cancer cell population almost reaches *x*_2_∼ 0.7 before the T-cell response is large enough to cause regression. In panel B we define the fundamental cycle period, *P*_*CIC*_, the time period between succesive peaks of the cancer cells (equivalently, the T-cell peaks). For larger values of *ϵ* (say *ϵ* = 0.9 shown in panel E), the T-cell response is triggered much earlier showing how the parameter regulates the relative timescales between these two populations. Figure 7 shows a panel of cancer-cell/T-cell interactions as the *θ* parameter varies from values *θ* = 1.0, 2.0, 3.0, 4.0, 5.0. For larger values of *θ*, the immune response controls the tumor cell population much more effectively, and the fundamental period decreases. As *ϵ* increases, the fundamental period decreases, following an approximate power-law formula *P*_*CIC*_ ∼ *αϵ*^−*β*^, shown in figure 8(a). Figure 8(b) shows the exponential scaling law governing the fundamental period as a function of *θ*.

**FIG 7.**
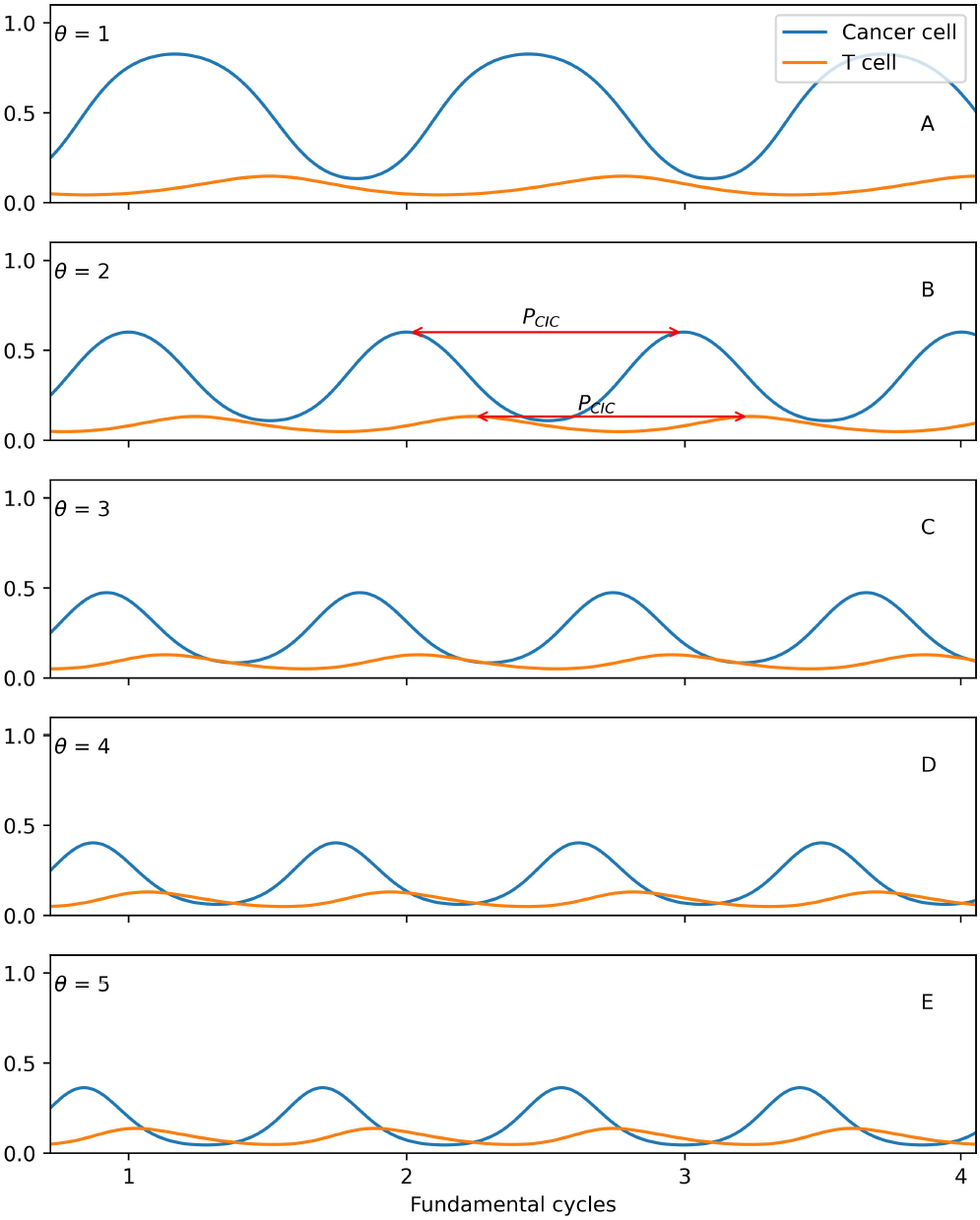
Dynamics of the system with respect to changes in the immunotherapy parameter *θ*, with *c* = 0.3. Time units are marked in cycle periods for *θ* = 2, which we take as the baseline setting for no immunotherapy. A. *θ* = 1; B. *θ* = 2; C. *θ* = 3; D. *θ* = 4; E. *θ* = 5.

**FIG 8.**
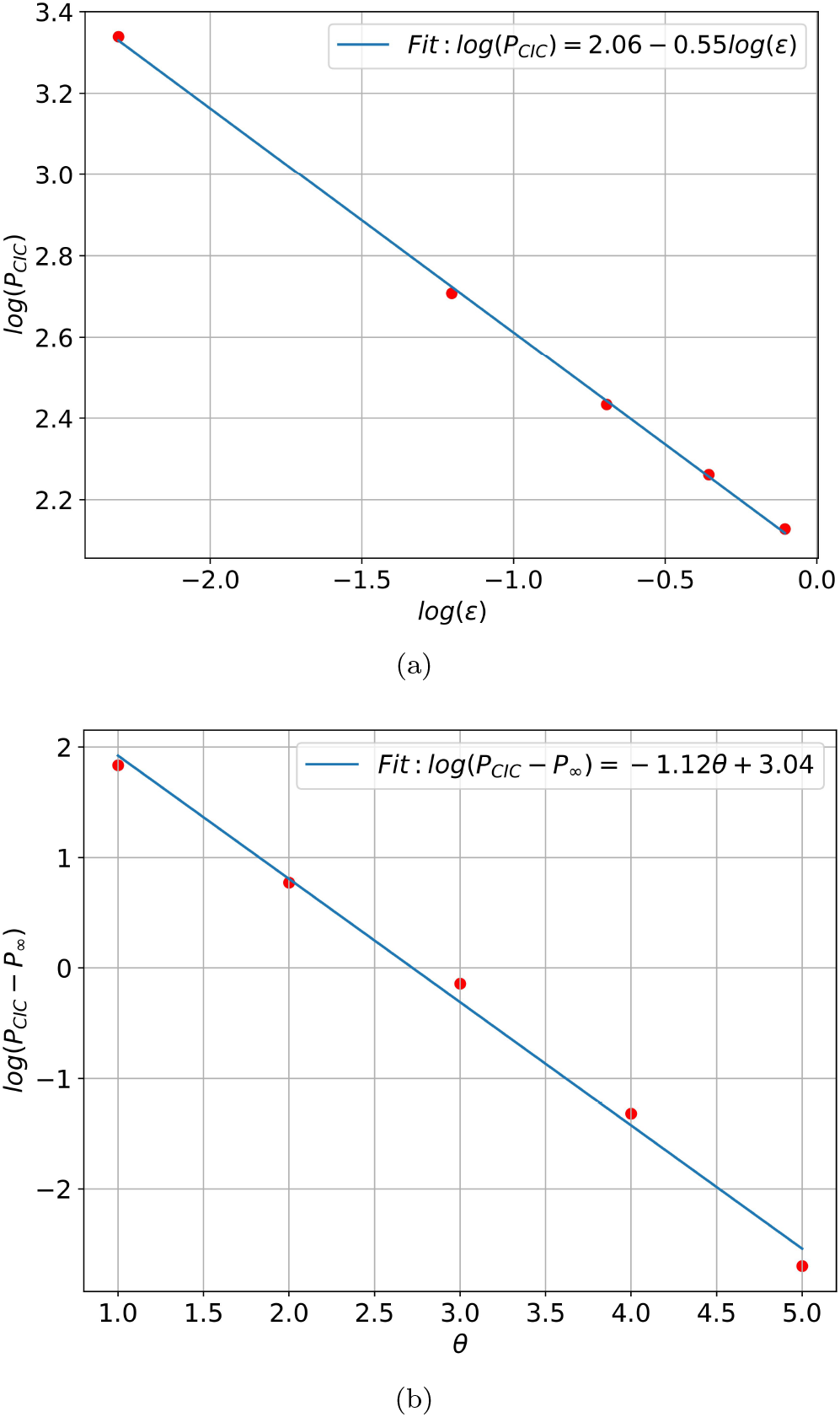
(a) Power-law scaling of the fundamental period with ϵ: *P* ∼*αc*^−*β*^; *α* ∼7.85, *β*∼ 0.55 and *θ* = 2. (b) Exponential scaling of the fundamental period with *θ*: (*P*− *P*_∞_) ∼ *α* exp (−*βθ*), *P*_∞_ = 12.83, *β* = 1.12, *α* = 20.91 and *c* = 0.3.

**FIG 9.**
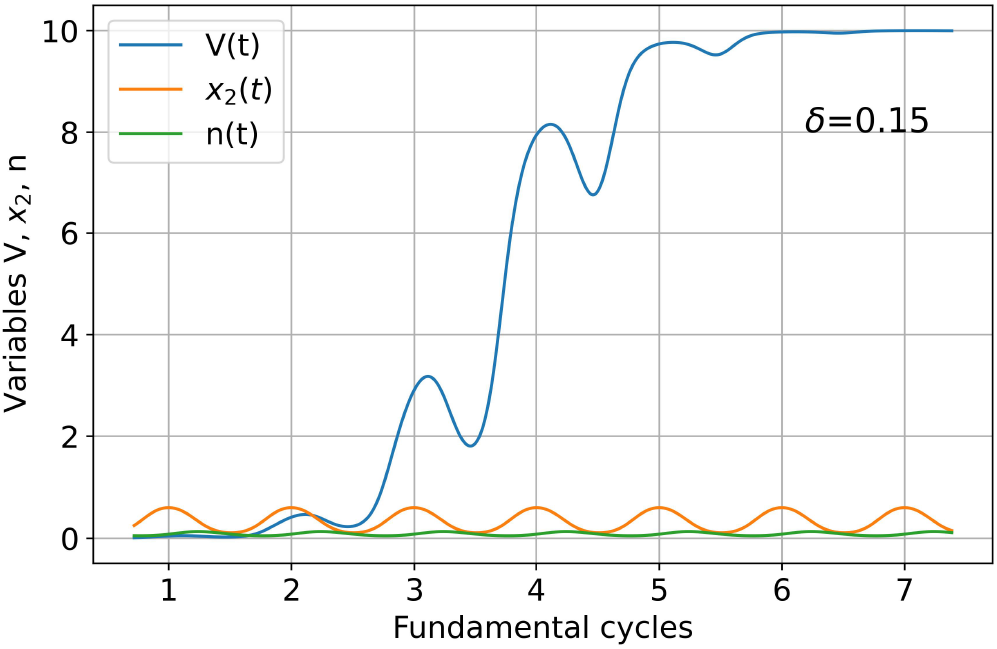
Volumetric growth *V* (*t*) through 6 fundamental cancer-immunity cycles. Shown are the curves *V* (*t*), and (*x*_2_(*t*), *n*(*t*)), with initial conditions *V* (0) = 0.01, *x*_2_(0) = 0.25, *n*(0) = 0.05, and carrying capacity *K* = 10, with *δ* = 0.15.

Using eqn (8) it is straightforward to derive a formula for the total volumetric growth or decay of the tumor volume *V* (*t*) through *m* fundamental periods *t* = *mP* :

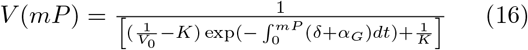

The first term in the denominator leads to overall exponential growth through each period *V* ∼ exp[*δmP*], modulated by the second term, 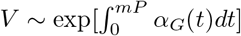 which could lead to growth or regression depending on the average sign of *α*_*G*_ over the cycle. Figure 9 shows a representative volumetric growth curve through six fundamental cancer-immunity cycles. Tumor regression due to the natural immune response is less and less pronounced as the tumor approaches its carrying capacity.

## III. DOSING SCHEDULES

We now show full simulations of the model system, eqns (1)-(3), (8) focusing on addressing the questions introduced in the beginning of the paper.

### A. Single-pulse dosing

We start by showing in figure 10 the dose-response curves associated with a single pulse of chemotherapy and single pulse immunotherapy, administered over a duration of 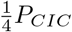.We compare the responses based on the timing of the pulses, first by starting the pulse at the peak of the tumor-cell curve (figure 10(a)), then starting at the trough (figure 10(b)). Based on the response of these pulses, it is clear that starting the chemotherapy pulse at the cancer cell peak, as the tumor is beginning to regress, and the immune response is beginning to rise, is more beneficial than starting at the trough. In figure 10(c) and (d) we show the dose-response curves using a single pulse of immunotherapy, also timed to begin at the peak (figure 10(c)), and then at the trough of the tumor-cell curve (figure 10(d)). In contrast to the timing of the chemotherapy pulse, for immunotherapy it is more beneficial to begin at the cancer cell trough, when the immune response is diminishing (cancer cell curve is increasing). In both the chemotherapy and the immunotherapy cases, our conclusion is that it is best to administer the respective pulses as the populations on which they act are in *decline*. Chemotherapy should be administered as the tumor-cell population begins to decline to accelerate the decline, immunotherapy should be administered as the immune response is in decline to stimulate a response. But since the tumor cell curves and the immune system curves are out of phase with each other (see figures 6 and 7), this means the pulses should also be out of phase.

**FIG 10.**
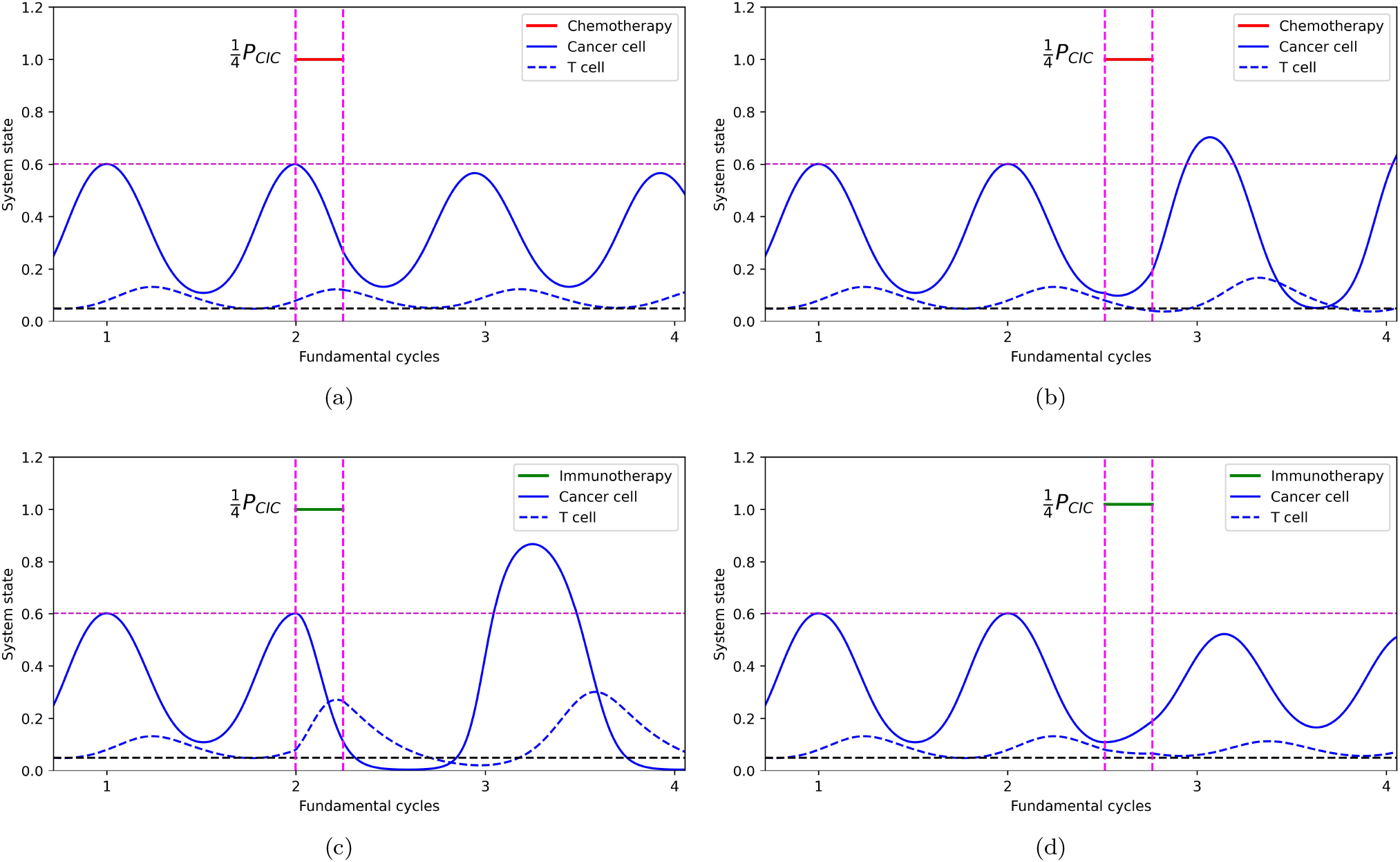
Single-pulse chemotherapy and immunotherapy dose response curves: *C* = 0 (off), *C* = 0.1 (on); *θ* = 2 (off), *θ* = 5 (on). Pulse is 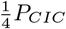 in duration. (a) Chemotherapy pulse beginning at cancer cell peak as immune system response rises and tumor begins a regression phase; (b) Chemotherapy pulse beginning at cancer cell trough as tumor begins a growth phase and immune system response is declining; (c) Immunotherapy pulse beginning at cancer cell peak as immune system response rises and tumor begins a regression phase; (d) Immunotherapy pulse beginning at cancer cell trough as tumor begins a growth phase and immune system response is declining.

Over what fraction of the cycle should the pulses act? This question is tied to the total dose, which is the area under the dose curve. In figure 11 we administer the single-pulse therapies over ^1^ *P*_*CIC*_, and through one full period. Figure 11(a) shows the dose-response for a pulse of chemotherapy through ^1^ *P*_*CIC*_, administered at the cancer-cell population peak, and 11(b) shows the response to immunotherapy administered at the trough. Figure 11(c) shows the dose-response for a pulse of chemotherapy through a full period, administered at the cancer-cell population peak, and 11(d) shows the response to immunotherapy administered at the trough. Comparing the responses of both with the results shown in figure 10 in which the duration was 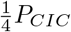,we have two important conclusions to draw. First, even though 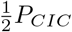 and full cycles of pulse chemotherapy deliver twice and four times the total dose as that shown in figure 10, the response is not as beneficial. *Careful timing can compensate for a lower dose*. Second, the chemotherapy pulse works best when administered through 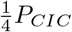 (e.g. figure 10(a)), whereas the immunotherapy pulse works best when administered through 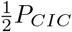 (e.g. figure 11(b)). In other words, the immunotherapy pulse should be sustained twice as long as the chemotherapy pulse. With this in mind, we next address the question of multi-pulse dosing schedules.

**FIG 11.**
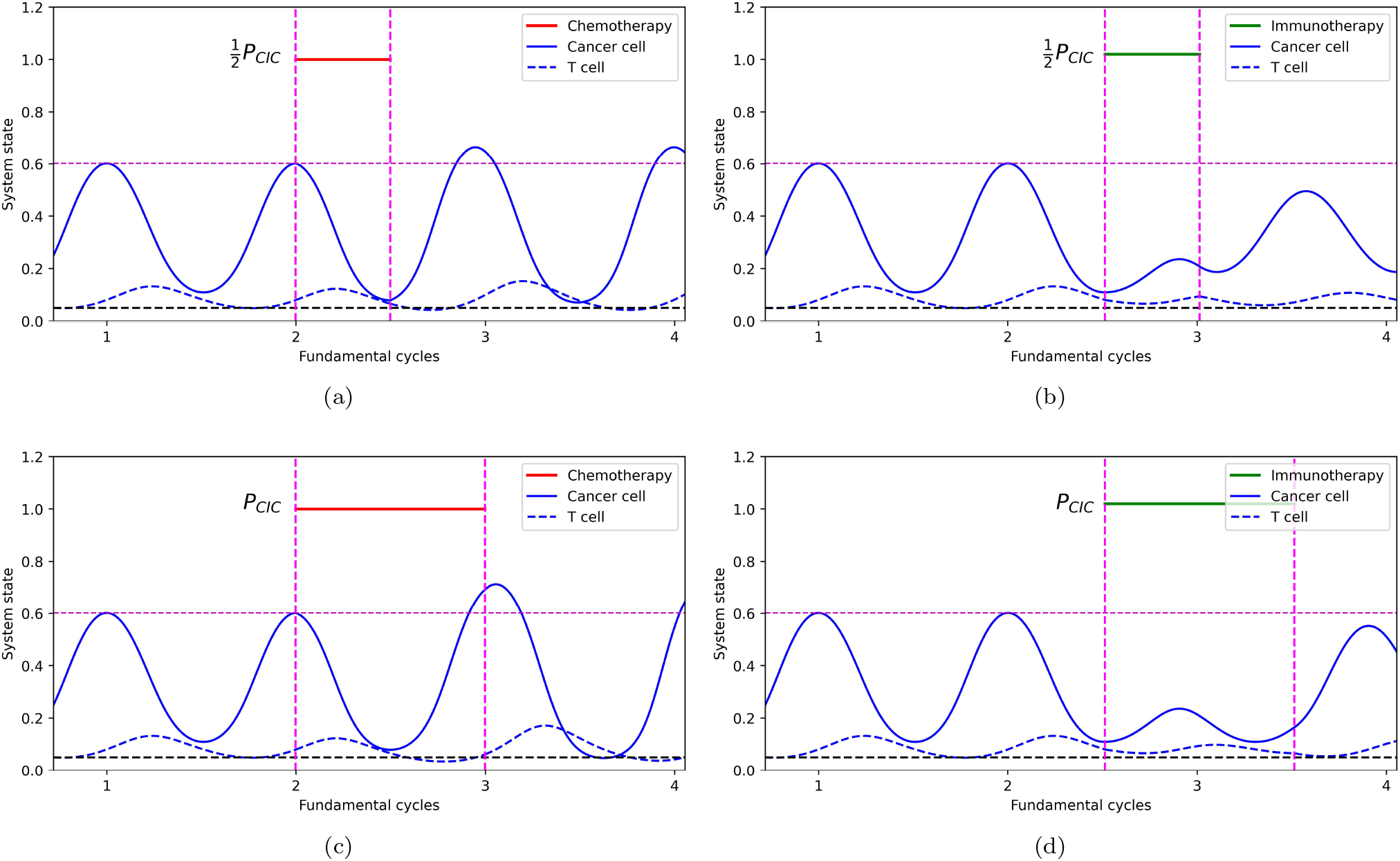
Single-pulse chemotherapy and immunotherapy dose response curves: *c* = 0 (off), *c* = 0.1 (on); *θ* = 2 (off), *θ* 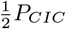 administered at the cancer cell trough; (c) Chemotherapy pulse is on for 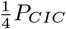 administered at the cancer cell peak; (d) Immunotherapy pulse is on for *P*_*CIC*_ administered at the cancer cell trough.

### B. Multi-pulse dosing

In figure 12 we use sequential multi-pulse dosing based on what we learned from the single-pulse dose responses. Figure 12(a) uses a chemotherapy dose of 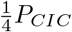 duration administered at the peak of the cancer cell curve, followed by an immunotherapy dose of 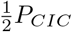 duration administered at the trough. Figure 12(b) shows the response when we reverse the ordering of the pulses. The results are non-transitive: reversing the order of the pulses leads to different outcomes, even if their size and duration are kept constant. From the response shown in figure 12, our conclusion is that preceding the chemotherapy pulse with a pulse of immunotherapy leads to a better response than if the ordering is reversed, as in figure 12(a).

**FIG 12.**
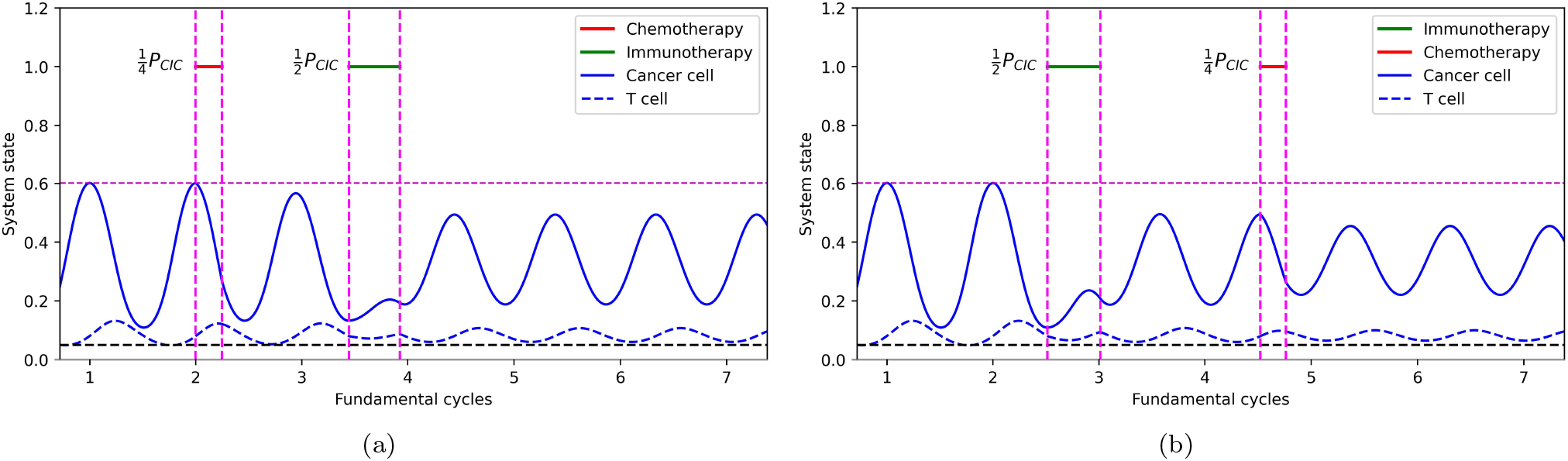
Multi-pulse sequential chemo-immunotherapy with chemotherapy beginning at the cancer cell peak and immunotherapy beginning at the trough. (a) 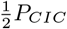 pulse of chemotherapy administered at peak, followed by 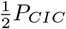 pulse of immunotherapy administered at trough; (b) 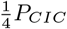 pulse of immunotherapy administered at trough followed by 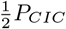 pulse of chemotherapy administered at peak. Note that the results are non-transitive. Immunotherapy pulse preceding the chemotherapy pulse is more beneficial.

In figure 13 we show responses to concomitant multipulse doses in which the chemotherapy and the immunotherapy doses begin simultaneously. In figure 13(a),(b) we administer the dosing for a duration of 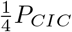,first at the cancer cell peak (figure 13(a)), then at the trough (figure 13(b)). For these two cases, administering the dosing concomitantly at the trough produces a superior outcome. However, figures 13(c),(d) show that for chemotherapy dosing administered for a duration of 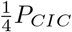 with immunotherapy dosing administered twice as long, for a duration of 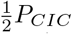,commencing at the trough of the cancer cell curve (figure 13(d)), yields the best results.

**FIG 13.**
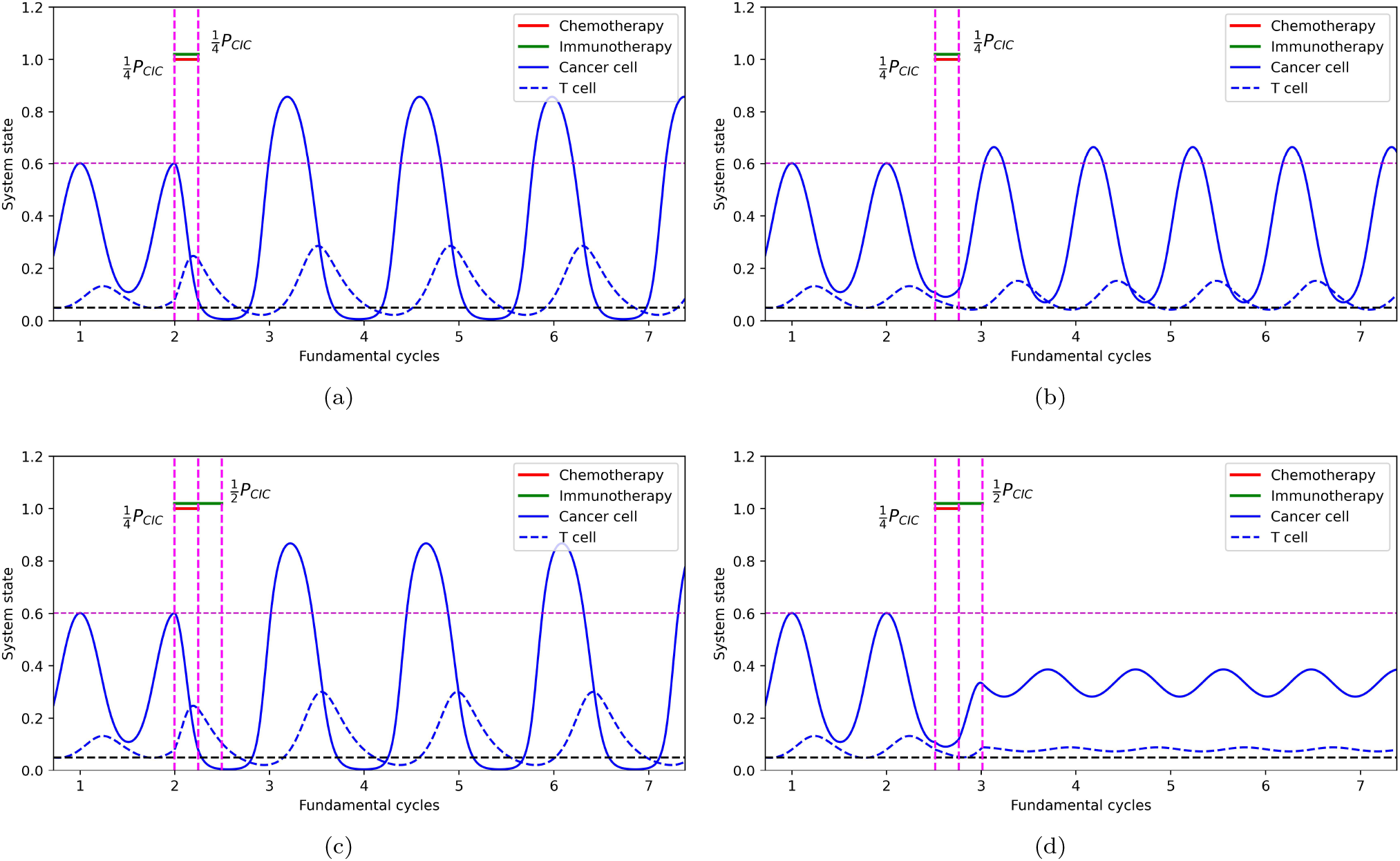
Multi-pulse concomitant chemo-immunotherapy administered at peaks and troughs. (a) 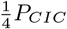 pulse of concomitant chemotherapy and immunotherapy pulses administered at peak; (b) 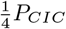 pulse of concomitant chemotherapy and immunotherapy pulses administered at trough; (c) 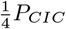 pulse of concomitant chemotherapy and 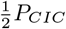 pulse of immunotherapy, both administered at peak; (d) 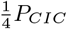 pulse of concomitant chemotherapy and 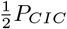 pulse of immunotherapy, both administered at trough is the most beneficial.

### C. Optimized dosing schedules over one fundamental period

We next turn to an optimal control treatment of the model. We would like to find an admissible controller which causes the system to follow a solution trajectory that minimizes a desired performance measure, or cost function:

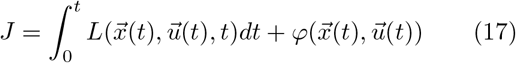

for controllers 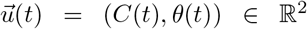 which are bounded above and below, *C*_*min*_ ≤ *C*(*t*) ≤ *C*_*max*_, *θ*_*min*_ ≤ *θ*(*t*) ≤ *θ*_*max*_, and with corresponding solution trajectories 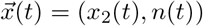. In particular, when we say that a controller is optimal, it indicates that

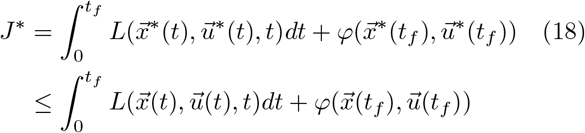

for all 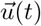 and corresponding solution trajectory 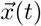 over a finite window of time 0≤ *t*≤ *t*_*f*_. The integral term is called the ‘running cost’, while the additive term is called the ‘terminal cost’.

We denote the optimal control schedule as 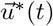 and the optimal trajectory as 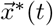.This inequality ensures that the optimal control and trajectory give us a global minimum of the performance measure *J*. We define the total dose delivered in time 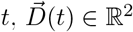,as

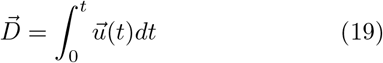

and thus,

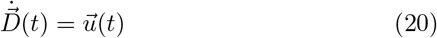

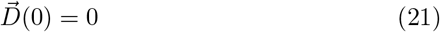

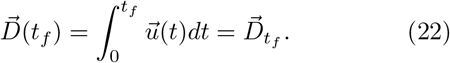

We keep a tight cap on the total dose delivered in our optimized solutions in order to best understand how to optimize the timing and dose levels within the restrictions of keeping toxicity low. In our case, we are only interested in minimizing the terminal value 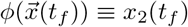 (*t*_*f*_ is the end of one fundamental period), which is called a classical Mayer problem (specifically choosing *L* = 0 and optimizing the terminal value) discussed in detail in [59]. The optimization problem becomes a two-point boundary value problem whose solution gives rise to the optimal trajectory 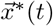 and the corresponding control 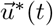.Optimal control for this problem was implemented using GEKKO, a Python package for machine learning and optimization of mixed-integer and differential algebraic equations. Documentation for this package can be found at https://gekko.readthedocs.io/en/latest/.

Figure 14 shows the results of our optimization over one fundamental cancer-immunity cycle (figure 14(a)), two half-cycles (figure 14(b)), and four quarter-cycles (figure 14(c)). All of the optimization windows work well as measured in terms of the final cancer-cell population fraction (i.e. the terminal cost), although optimizing once over a full cycle, as shown in figure 14(a) seems to produce the best outcome. We also point out that the optimized dosing schedules in this case are relatively simple, and follow the same general two principles of the multi-pulse schedules: (i) immunotherapy dosing is administered (roughly) twice as long as chemotherapy dosing, and (ii) immunotherapy dosing precedes the chemotherapy dosing. Notice that we start our optimization window at the peak of the cancer cell curve and end at the next peak, so a simple statement about the optimal starting times, as in the multi-pulse case, is not as clear. This is because we are fixing the start and end times.

**FIG 14.**
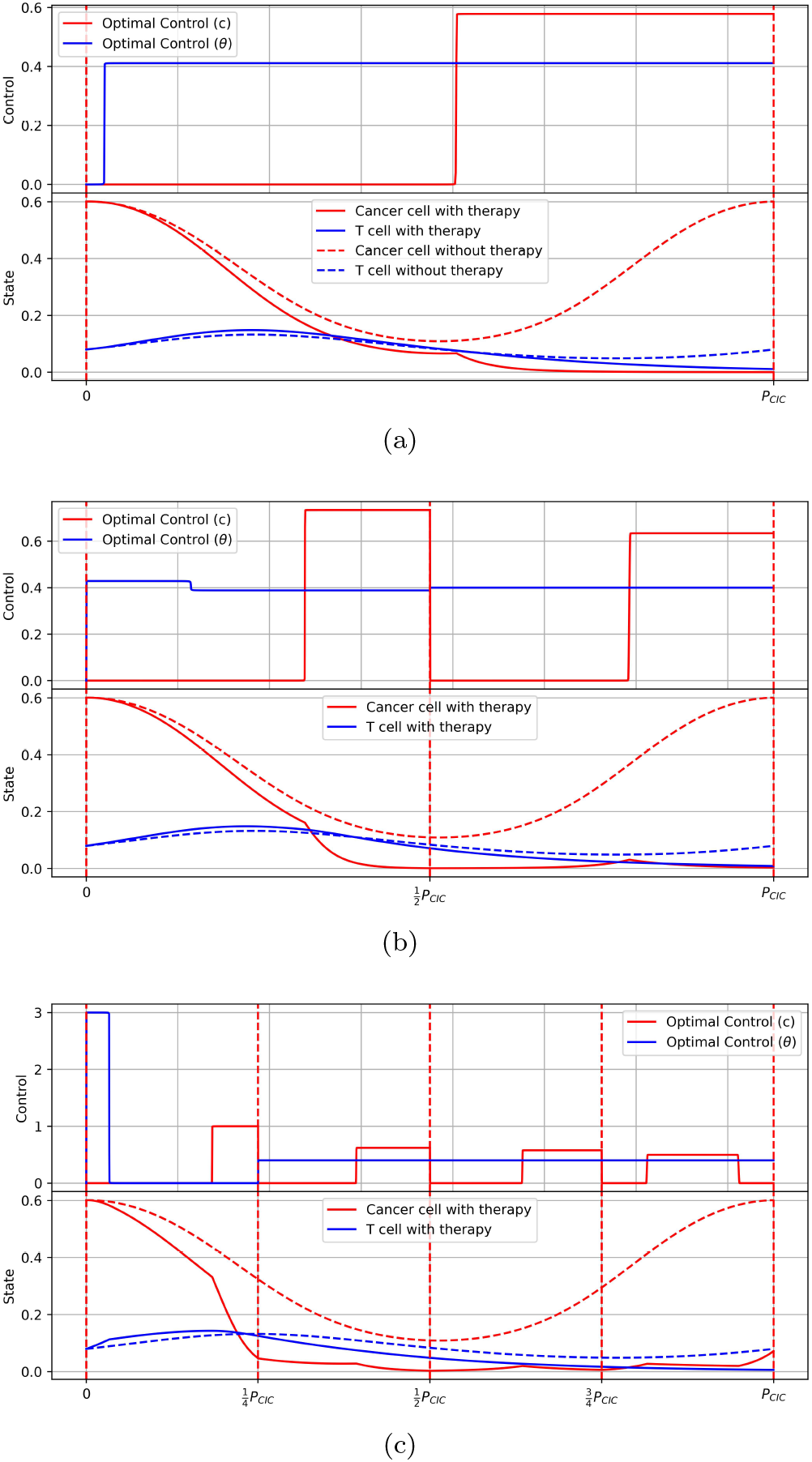
Optimized chemo-immuno therapy dosing schedules which minimize the final cancer cell fraction at the end of one fundamental period (15.0 time units). Shown in each panel are the dosing schedules and the controlled and uncontrolled cancer cell and T-cell fractions 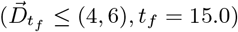.The *y*-axis for the Control panels shows the actual *C* value used (when on). For the *θ* value, add 2 to the *y*-axis value (since 2≤ *θ* ≤ 5). (a) Optimized dosing schedules and responses where optimization is performed through the full period; (b) Optimized dosing schedules and responses where optimization is performed through two half-periods; (c) Optimized dosing schedules and responses where optimization is performed through four quarter-periods. Notice the rise in the cancer cell fraction at the end due to the fact that the total chemothreshold was reached.

**FIG 15.**
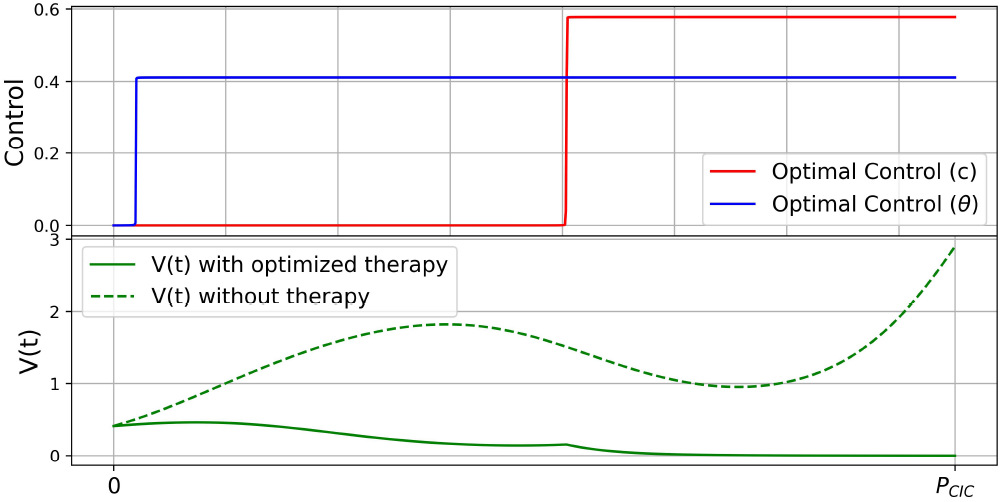
Volumetric growth *V* (*t*) using the full cycle optimization shown in figure 14(a). Shown are the optimized chemotherapy and immunotherapy schedules (above), and *V* (*t*) with and without therapy (below).

## IV DISCUSSION

The cancer-immunity cycle begins with the presence of a cancer cell that has accumulated the appropriate genetic alterations endowing it with an evolutionary fitness advantage over the surrounding healthy cell population in which it competes for resources. The subsequent growth of the cancer cell population at the expense of the healthy cells is captured in the first term, *A*_*G*_, of the payoff matrix (9) which is of prisoner’s dilemma type [50]. As the cancer cells proliferate, they exploit their fitness advantage in what is often called a tragedy of the commons [50] -the cancer cells saturate the tumor as an evolutionary stable state (ESS), but the overall average fitness of the healthy/cancer cell mix declines. The cancer cell’s surface over-expresses an antigen marking it as a potential target for attack by the surveilling T-cell population which deliver appropriate antibodies to kill the cell. The increase in antigen level triggers an increase in the T-cell population which begin to locate, infiltrate, and kill the tumor cells. This is captured by the growth equation (3) for *n*(*t*) whose growth rate becomes positive in the presence of a sufficiently high proportion of cancer cells (i.e. antigen levels).

As the T-cell population grows, the linear coupling term (10) shifts the balance of terms in the payoff matrix towards the second term *A*_*R*_, triggering tumor regression instead of growth. As the cancer cell population decreases, the T-cell population declines, again due to the coupling term in (4). When this population declines further, the ability of the T-cells to locate and deliver the antibodies needed to kill the cancer cells diminishes to the point when the first term in the payoff matrix *A*_*G*_ again dominates, and the cycle begins anew. This oscillatory interchange establishes the fundamental period (and frequency) of the baseline cancer-immunity cycle. The framework highlighted in figure 1, also includes the additional roles of our chemotherapy dosing function *C*(*t*) which directly selects out the cancer cells (sometimes leading to chemoresistance [37, 54]), and our immunotherapy dosing function *θ*(*t*) (eqn (4)) which triggers the growth term of the T-cell equation, thereby indirectly helping to select out the cancer cells. Both of these dosing schedules are chosen to synchronize with the underlying period of the cancer-immunity cycle.

Our simplified model captures several key features of the more complex and detailed cancer-immunity cycle [19], already a simplification of the true process, framing it as a evolutionary game played between the cancer cell, healthy cells, and T-cells. As such, it is well suited to address questions associated with timing and sequencing of chemotherapy and immunotherapy schedules delivered to populations of cells competing under selection dynamics. There are, of course, certain potentially important biological mechanisms it leaves out. For example, it does not explicitly include a mechanism for the build-up of cancer antigens in the blood as the cycle repeats [11, 19]. Since the model is deterministic, it does not include any effects of stochasticity [55, 60], potentially important in the early stages of tumor growth where the cancer cell population level is low. It does not include any mechanisms for the evolution of chemo-resistance [14, 15, 61], or any ability to model different mechanisms and modes of immunotherapy [14, 15, 18] or chemotherapy. Furthermore, it is likely that there is variablility in the details of the cancer-immunity cycle across patients — this is not yet well enough understood to try to optimize and exploit.

The model can be developed further in the future to include some of these effects, specifically by adding in more details associated with the TME subcycle [11], more subpopulations competing in the evolutionary game (leading to a higher dimensional payoff matrix), and more compartments to delineate types of immune cells and their various roles as the cycle unfolds. But all of these additional features will come at the expense of the relative simplicity and clarity the current model provides.

## V. CONCLUSION

Since the introduction of the cancer-immunity cycle framework in 2013 [19], and its subsequent recent update in 2023 [11], it has provided a very unifying way of organizing the many complex biological processes controlling the interaction of the tumor and the immune system. Despite this, as far as we are matical models that have been introduced to capture its features, and used to build on.

While the mathematization of a complex set of biological events cannot, in and of itself, supply a mechanistic understanding of the events, it can provide a useful framework in which to test and explore hypotheses associated with them, one of which is the important role of timing and synchronization of chemotherapy and immunotherapy in an evolutionary environment touched on in [47]. From a control theory and nonlinear dynamical systems perspective, it is certainly not surprising that using time-dependent controllers (each with their own characteristic timescales) to shape the trajectories of a nonlinear evolutionary system (with its own inherent timescales) is a delicate and tricky undertaking with the ability to create many different outcomes even with small amplitude controllers. Small changes in these exogenous control functions can produce large and sometimes important changes in the endogenous state-variable trajectories if synchronized appropriately. In an engineering/physics setting, this fact is widely appreciated[32] and generally referred to as suppling a forcing function that is tuned to a resonant frequency of the underlying system. In biological settings, since complex feedback/adaptive control protocols are much more challenging to implement due to limited ability to use sensors and monitors in stochastic in vivo settings [62], there is less of an appreciation of the vast range of new possibilities that time-dependent controllers, if synched optimally with the underlying dynamical system, can potentially provide.

Two general conclusions based on our model, that strategic dose timing can compensate for a lower total dose, and that chemotherapy schedules and immunotherapy schedules should each synchronize with the phases of the respective populations on which they act, open up many exciting new avenues for less toxic and more effective multi-modal dosing strategies that seek to exploit tumor evolution as a potentially effective therapeutic target [49].

## ACKNOWLEDGMENTS

We gratefully acknowledge support from the Army Research Office MURI Award #W911NF1910269 (2019-2024). PKN also gratefully acknowledges the support of the Guggenheim Foundation through their 2024 Fellows Program (Applied Mathematics).

